# The single-species metagenome: subtyping *Staphylococcus aureus* core genome sequences from shotgun metagenomic data

**DOI:** 10.1101/030692

**Authors:** Sandeep J. Joseph, Ben Li, Robert A. Petit, Zhaohui S. Qin, Lyndsey A. Darrow, Timothy D. Read

**Affiliations:** Department of Medicine, Division of Infectious Diseases Emory University School of Medicine, Atlanta, Georgia, USA; Department of Human Genetics Emory University School of Medicine, Atlanta, Georgia, USA; Department of Biostatistics and Bioinformatics Rollins School of Public Health, Emory University, Atlanta, Georgia, USA.; Department of Epidemiology Rollins School of Public Health, Emory University, Atlanta, Georgia, USA.

## Abstract

Metagenome shotgun sequence projects offer the potential for large scale biogeographic analysis of microbial species. In this project we developed a method for detecting 33 common subtypes of the pathogenic bacterium *Staphylococcus aureus.* We used a binomial mixture model implemented in the *binstrain* software and the coverage counts at > 100,000 known *S. aureus* SNP (single nucleotide polymorphism) sites derived from prior comparative genomic analysis to estimate the proportion of each subtype in metagenome samples. Using this pipeline we were able to obtain > 87% sensitivity and > 94% specificity when testing on low genome coverage samples of diverse *S. aureus* strains (0.025X). We found that 321 and 149 metagenome samples from the Human Microbiome Project and metaSUB analysis of the New York City subway, respectively, contained *S. aureus* at genome coverage > 0.025. In both projects, CC8 and CC30 were the most common *S. aureus* subtypes encountered. We found evidence that the subtype composition at different body sites of the same individual were more similar than random sampling and more limited evidence that certain body sites were enriched for particular subtypes. One surprising finding was the apparent high frequency of CC398, a lineage associated with livestock, in samples from the tongue dorsum. Epidemiologic analysis of the HMP subject population suggested that high BMI (body mass index) and health insurance are risk factors for *S. aureus* but there was limited power to find factors linked to carriage of even the most common subtype. In the NYC subway data, we found a small signal of geographic distance affecting subtype clustering but other unknown factors influence taxonomic distribution of the species around the city. We argue that pathogen detection in metagenome samples requires the use of subtypes based on whole species population genomic analysis rather than using ad hoc collections of reference strains.

## Introduction

Bacterial species are commonly comprised of multiple phylogenetic clades that have distinctive phenotypic properties. The process of identifying which clade a bacterial strain belongs in goes by several names but here we will refer to it as *subtyping.* Commonly used subtyping methods include multilocus sequence typing (MLST), pulsed-field gel electrophoresis (PFGE), oligotyping and variable-number of tandem-repeat typing (VNTR) [1], Each of these these methods was developed for bacteria first isolated in pure culture in the laboratory before DNA extraction. For early disease diagnosis of pathogenic bacterial species and to understand bacteria in the context of their natural community, it would be advantageous to subtype directly from clinical specimens such as blood and sputum. However, current direct identification options such as 16S rRNA gene sequence, FISH and REP-PCR, are not able to subtype bacteria to below the species level taxonomic resolution, nor to deal with mixtures of subtypes of the same species being present.

Over the past few years, the expansion of ‘metagenomic’ culture-free shotgun sequencing of clinical and environmental DNA, exemplified by the human microbiome project (HMP), has generated a plethora of revolutionary new analysis methods. This includes a set of tools that has been developed to classify the abundance of bacteria to the taxonomic level of species or genera in metagenomic data sets (see Discussion). Within metagenomic data sets there are often hundreds to thousands of individual reads from the more abundant bacterial species, potentially providing enough evidence for subtyping [2], Here we report the development of a preliminary metagenome based subtyping scheme for *S. aureus* and the results obtained when we probed for the distribution of common clades across human body sites and the built environment.

*S. aureus* is one of the most common hospital infections, often causing chronic diseases with poor outcome [3][4][5][6] [7] [8] [9], *S. aureus* is also a problem outside the hospital as a community-acquired bacterial infections in humans [10] [11], livestock [12] [13] and other animals[14], *S. aureus* is a common asymptomatic colonizer of humans, with the nares (nose) believed to be the most important site. Estimates for human nasal carriage rates suggest ~20-50% of humans are persistently colonized with *S. aureus*, with 60-100% of individuals harboring *S. aureus* at some point in their lifespan [15][16,17], The population of *S. aureus* asymptomatically colonizing the nose in healthy individuals is thought to be a major source for transmission [18]. Many studies have shown that persistent nasal carriage of *S. aureus* is a risk factor for pathogenic infection, but the overall association of these carriage strains to the presence of endogenous strains that establish pathogenic infections is currently unknown [15] [19] [20].

*S. aureus* strains can be classified into a limited number of clonal lineages of related MLST sequence types (clonal complexes[21]), which differ in their geographical distribution and propensity to cause human diseases. The acquisition of the SCCmec cassette, producing the MRSA strains is more common in some clonal lineages than others[22], as is the acquisition of *vanA* genes to produce VRSA (vancomycin resistant *S. aureus*)[23], It is important to understand the genetic diversity (population structure) of *S. aureus* strains that colonize the different body sites in order to understand how commensal strains present in healthy human population might act as a predisposing factor for future invasive infections. Using our subtyping scheme based on metagenome data, we performed epidemiological modelling to understand whether there is any association with the demographic and life history characteristics collected using the responses that subjects gave to an extensive survey, and the different strains of *S. aureus* identified at each body site.

## Results

### An *S. aureus* subtyping scheme for metagenomic data

There were two main challenges in subtyping a bacterial species based on metagenomic data: missing loci and mixed strain composition. The first problem arises when coverage is too low to guarantee obtaining sequence across the entire genome. This obviates against using schema that have an absolute requirement for a particular sequence being represented, such as VNTR and MLST. The mixed strain issue meant that the scheme cannot assume that only one subtype is present. For these reasons we chose to use a software, *binstrain[24]* that we had previously developed for distinguishing mixed clonal populations of *Chlamydia trachomatis.* The *binstrain* approach used a binomial mixture model to separate the relative abundance of subtype-specific SNPs (other types of variants can be used but in this report we are focussing only on SNPs) found across the population of sequence reads. The software, implemented in R, integrated the results from SNP positions found across the whole or partial genome sequence and was not dependent on specific loci. Further, since *binstrain* was developed to determine sample mixtures, results were expressed as the likely proportion of each subtype present in the sample. *binstrain* required as input a matrix of SNPs assigned to each subtype. To generate this matrix we used an iterative approach to sample the number of definable subtypes based on currently available *S. aureus* genome sequence data. We first estimated that there were 19 major *S. aureus* population groups in 43 completed genomes aligned using the MAUVE program[25] by performing fineSTRUCTURE population subdivision analysis[26] on the the core portion of chromosomes (Supplementary Table 1). We then randomly picked one genome from each of the subtypes as a representative and created a matrix of each SNP position relative to the inferred ancestral nucleotide. We named the subtypes based on the clonal complex (CC) that contained most strains. This subtyping scheme largely overlaps the *S. aureus* MLST scheme (in terms of assigning strains in individual sequence types to larger clonal complexes[27]. We generated an inferred ancestral sequence (SA_ASR) based on the MAUVE alignment and tree and used it as a reference for alignment of sequence reads from FASTQ data. The reason for using the SA_ASR was to attempt to reduce the possible ascertained effect of alignment bias in closely related strains that could occur if a modern reference was chosen instead.

We tested the performance of *binstrain* with the 19 strain SNP matrix to classify the raw sequence of 2,692 genetically diverse *S. aureus* strains downloaded from the Short Read Archive database. Each strain was assigned to a MLST sequence type (ST) based on the results of the SRST software [28]. The input to *binstrain* in each case was the SNP matrix and an output generated by the samtools mpileup command of the raw metagenomic reads matched against the SA_ASR inferred ancestral chromosome sequence. We reported a positive result when the highest binstrain ***β*** value was above the threshold of 0.8 (an arbitrarily chosen value). We found the sensitivity of the initial subtyping scheme was too low, with 1,379 genomes not assigned to any subtype, a reflection of the sampling of the *S. aureus* diversity in the current stock of completed genomes. To allay this problem, we selected 21 unfinished genome projects from diverse CCs not represented in the original set. We repeated the previous steps of creating an alignment of the core conserved chromosome, inferring ancestral states and re-constructing a SNP matrix. The forty representative genomes contained a total of 102,057 SNP positions in 2,872,915 bps of the *S. aureus* chromosome.

Before attempting to subtype *S. aureus* data from the HMP metagenomic project, we ran experiments on simulated data (100 bp Illumina reads generated by the ART program[29]) to test the sensitivity and specificity of the method. First, we showed that the presence of a large excess of sequence reads from the near-neighbor species *S. epidermidis (S. epidermidis ATCC12228* (accession number: NC_004461.1) and *S. epidermidis RP62A* (accession number: NC_002976.3) did not affect the accuracy of the binstrain assignment. This was an expected result as the *S. epidermidis* genome contained very few SNPs that overlapped those used by *binstrain* for subtype identification and therefore is effectively ‘invisible’ to this typing method. We next created 3 artificial nasal metagenome consisting of reads from the abundant PATRIC pathogens in the human microbiome that have been well characterised in the HMP population [30] (common nasal microflora), *Propionibacterium acnes, Staphylococcus epidermidis*, *Streptococcus mitis*, *Bacteroides vulgatus, Staphylococcus haemolyticus,* and *Staphylococcus saprophyticus,* and included synthetic reads from 3 randomly chosen *S. aureus* genomes that had a combined length of ~2.9 Mb (ie ~ 1X genome coverage). Despite being less less 0.5% of the total reads in the artificial metagenome, we were able to accurately determine the approximate relative proportions of the *S. aureus* subtypes (Figure 1). In each tests 9-14% of the variation was attributed to a range of subtypes other than the three used in creation of the synthetic microbiomes (marked as “other’ in Figure 1). We believe these miscalls come from a combination of sequence error introduced by the ART software and binstrain assignment of probability to multiple subtypes to SNPs in internal branches of the species phylogeny.

**Figure 1.**
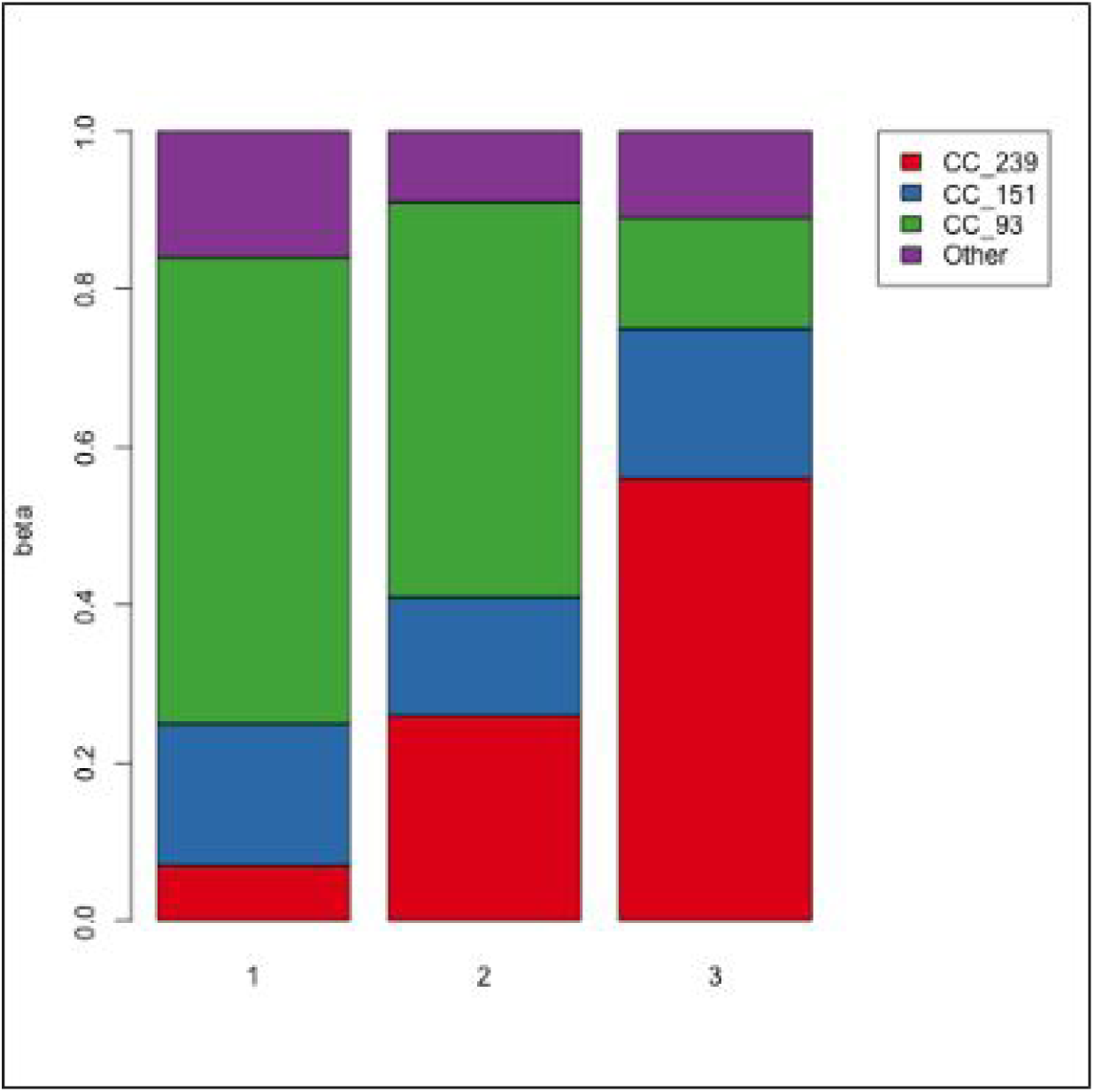
Binstrain results from synthetic microbiomes. *S. aureus* reference genomes (CC_151, CC_239 and MLST_93) were added to *Propionibacterium acnes* (200X), S. epidermidis (100X), *Streptococcus mitis* (7X), *Bacteroides vulgatus* (5X), *Staphylococcus haemolyticus* (4X) and *Staphylococcus saprophyticus* (3X). **1) microbiome 1** – CC_151_151 (0.5X), CC_239_239 (0.1X) and MLST_93 (0.25X) **2) microbiome 2** – CC_151_151 (1.5X), CC_239 (1X) and MLST_93 (0.75X) and **3) microbiome 3** – CC_151_151 (1.0X), CC_239 (5.0X) and MLST_93 (2.5X). Barplots show the results from binstrain analysis. The ‘other’ category represent diverse subtypes called with low probability.

To further assess the sensitivity and specificity of the *binstrain* test, we performed analysis on 10 simulated read sets of between 0.00625X and 5X genome coverage for each of the 40 subtype-defining genomes and repeated the analysis 9 times (a total of 3600 FASTQ files and *binstrain* runs). The binstrain test showed a low false positive rate with the simulated data: we only found strains assigned to the wrong subtype on 748/3600 times (20.77%), and this only occurred when the coverage was low (< 0.0125X coverage) and the misclassification was to between subtypes within the same CC5 (Figure 2). The false negative rate was high at the lowest coverages, but at 0.5X coverage and above > 95% of strains were correctly assigned. Overall the results showed a bimodal distribution (Figure 2 and Supplementary Figure 1), tending to produce either a **β** < 0.1 or > 0.8 with few points in between.

**Figure 2.**
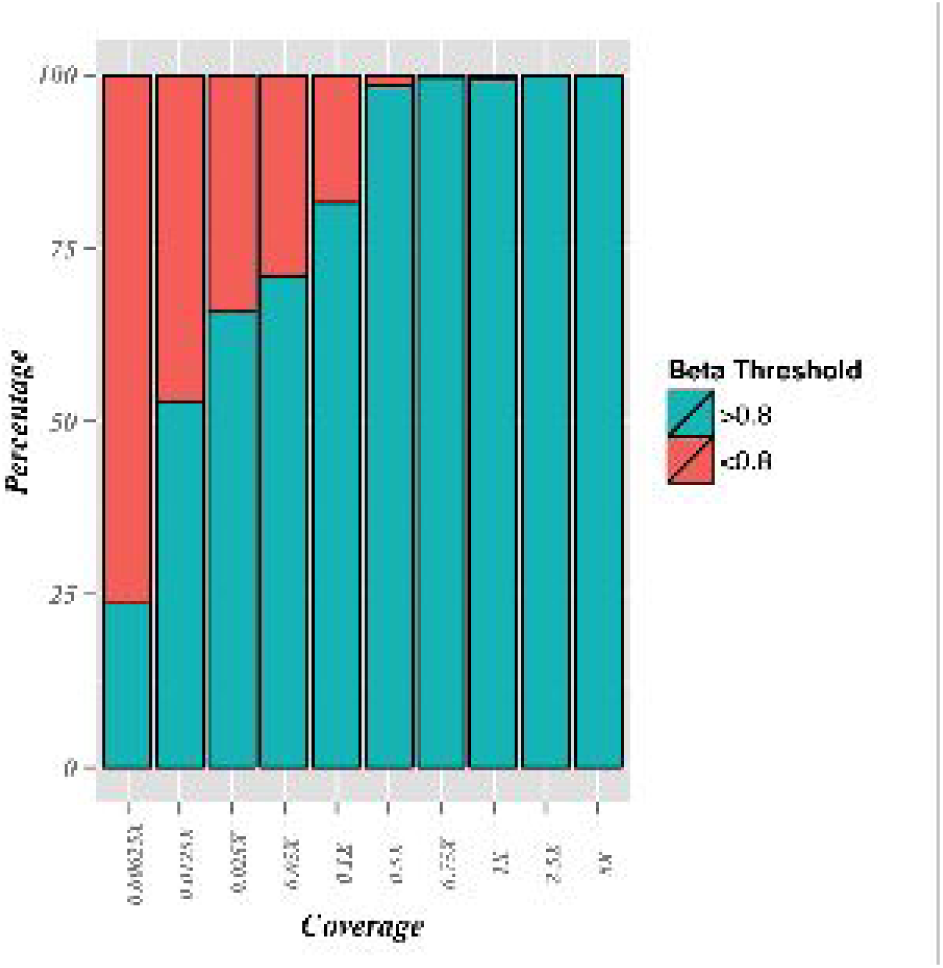
Sensitivity of binstrain assignment at different coverages. Synthetic FASTQ files of 40 strains were at converage genome coverages were challenged. Plots show percent correctly called above beta > 0.8 threshold.

As a final test, we reran the second version of our SNP matrix against 2,114 *S. aureus* FASTQ files downloaded from the NCBI SRA, representing a random selection of genotypes. For each strain we determined the true MLST type using the SRST tool [28] on 50X genome coverage data. Each FASTQ file was then randomly downsampled to subsets of 0.025X - 0.75X coverage and run through the *binstrain* classifier (Tables 1a and b Supplementary Table 2).

**Table 1(a).** Sensitivity of the v2 subtyping matrix on 2114 *S. aureus* FASTQ files from the NCBI SRA at different cutoff **β** values

| Coverage | $\beta \geq 0.90$ | $\beta \geq 0.80$ | $\beta \geq 0.70$ | $\beta \geq 0.60$ |
| --- | --- | --- | --- | --- |
| 0.025X | 83.91% | 86.19% | 86.80% | 87.41% |
| 0.05X | 87.04% | 89.03% | 89.64% | 89.97% |
| 0.1X | 88.70% | 91.77% | 92.33% | 92.76% |
| 0.5X | 85.71% | 93.18% | 95.93% | 97.20% |
| 0.75X | 84.34% | 92.38% | 95.78% | 97.25% |

**Table 1(b).** Specificity of the v2 subtyping matrix on 2114 *S. aureus* FASTQ files from the NCBI SRA

| Coverage | $\geq 0.90$ | $\geq 0.80$ | $\geq 0.70$ | $\geq 0.60$ |
| --- | --- | --- | --- | --- |
| 0.025X | 91.69% | 93.94% | 94.54% | 94.98% |
| 0.05X | 94.70% | 96.45% | 97.05% | 97.21% |
| 0.1X | 94.37% | 97.17% | 97.46% | 97.46% |
| 0.5X | 90.65% | 95.53% | 96.07% | 96.46% |
| 0.75X | 90.26% | 96.69% | 97.53% | 98.25% |

Compared to the synthetic FASTQ files reported above we found that the sensitivity of the test was higher with data sets from real sequencing projects ( ~84% on 0.025X coverage data even with a restrictive **β** cutoff of 0.9 or above (Table 1a)). When the positive calls were mapped onto a whole genome phylogeny of the *S. aureus* isolates we confirmed that there were no major lineages systematically excluded (Figure 3). Some individual strains with long branches connected to the rest of the tree were missed. Treating these strains as individual representatives of novel subtypes may improve accuracy of the next iteration of the *S. aureus* scheme.

**Figure 3.**
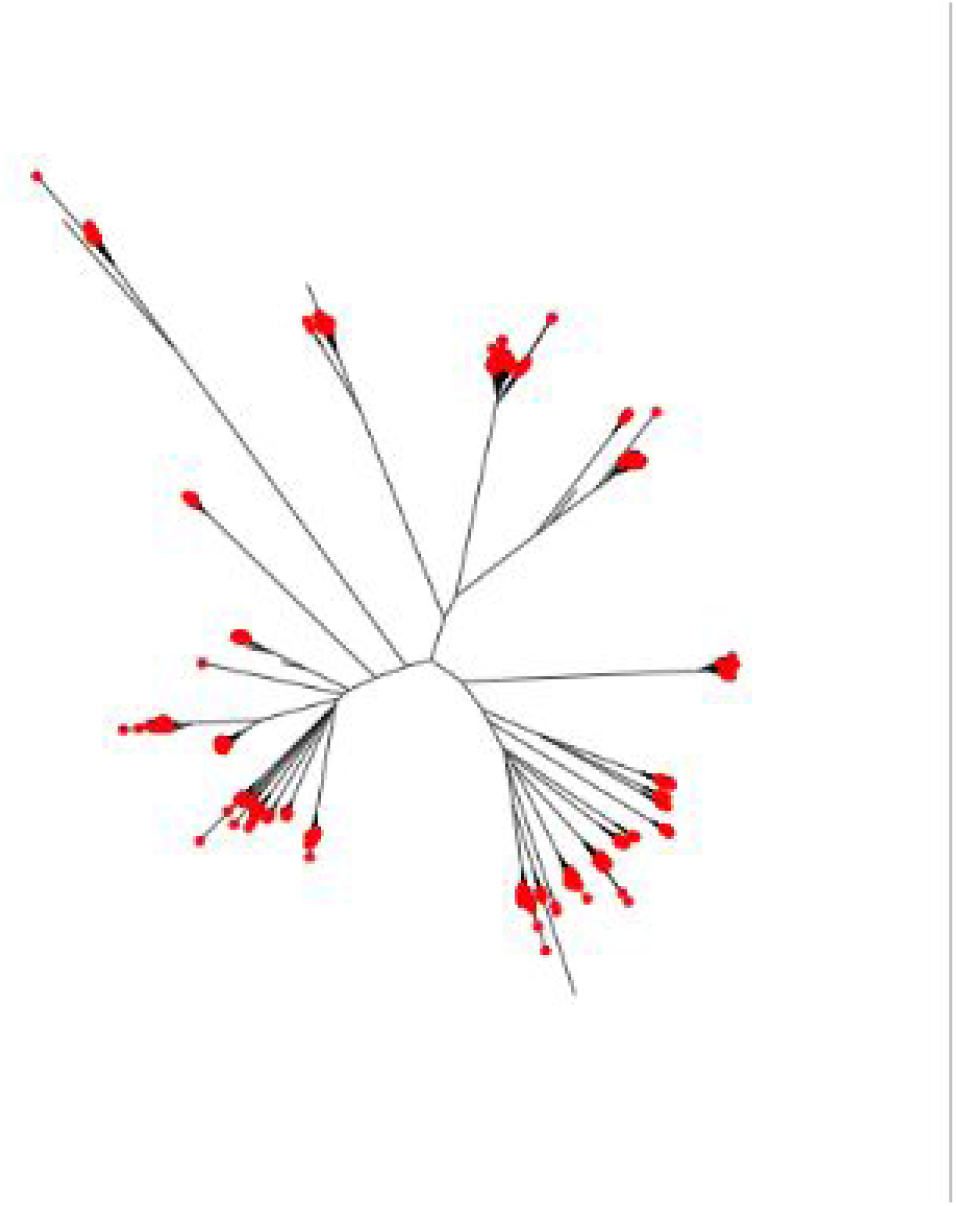
Phylogenetic distribution of *S, aureus* strains typed with beta > 0.65. Each strain classified into a subtype is marked with a red point. The neighbor-joining, tree was produced from a distance matrix derived from concatenation of all variable positions mapped against reference strain N315.

We measured specificity using the logic that strains with the same ST should be placed in the same subtype; thus we counted the largest subtype for each ST as ‘true’ (Table 1b). CC8, CC5 and CC30 were originally partitioned into multiple subtypes but in analyzing these results we noted considerable cross-specificity. Therefore we decided to collapse each clonal complex into only one subtype each, resulting in a final total of 33 subtypes. Based on these criterion we achieved > 91% specificity even at the lowest coverage and most stringent **β** cutoff. We noted that strains classified as ST5 had a lower specificity than other clonal complexes (Supplementary Table 2). In fact, 97/157 ST5 strains were only classified as CC5 and the rest were grouped as CC15, CC97 and CC8. At this time we do not know the reason for this misclassification.

**Table 2.** *S. aureus* positive HMP body sites based on reads mapping to the SA_ASR sequence. Percentages based on total number of samples for that body site.

| Body site | Total number of | Number of *S.* | Number samples with |
| --- | --- | --- | --- |

|  | **samples** | ***aureus* positive samples** | **> 0.025X *S. aureus* coverage** |
| --- | --- | --- | --- |
| Anterior nares | 137 | 78(57%) | 68(50%) |
| Attached keratinized Gingiva | 14 | 4(29%) | 4(29%) |
| Buccal mucosa | 185 | 56(31%) | 56(30%) |
| Hard Palate | 1 | 1(100%) | 1(100%) |
| Left retroauricular crease | 23 | 23(100%) | 23(100%) |
| Palatine tonsils | 19 | 7(37%) | 6(32%) |
| Posterior fornix | 108 | 11(10.20%) | 9() |
| Right Antecubital fossa | 1 | 1(100%) | 1(100%) |
| Right retroauricular crease | 31 | 28(90%) | 28(90%) |
| Saliva | 7 | 2(28.57%) | 1(14%) |
| Stool | 251 | 14(6%) | 7(0.3%) |
| Subgingival plaque | 19 | 1(5%) | 1(5%) |
| Supragingival plaque | 210 | 65(31%) | 37(18%) |
| Tongue dorsum | 221 | 82(37%) | 82(37%) |

Based on these tests, we decided that a coverage of 0.025X *S. aureus* genome equivalents was a reasonable threshold for our HMP metagenome data, giving us a large sample set with acceptable specificity and sensitivity errors of ~ 11% and 8%, respectively. We performed the same analysis using a 0.5X cutoff and saw similar patterns to that when the 0.025X threshold was used.

### Assignment of *S. aureus* subtypes in the HMP metagenomic dataset using whole genome subtyping

Having developed and tested the *S. aureus* binstrain classifier we attempted to call the subtypes present in 1,263 whole metagenomic sequencing samples from the healthy human cohort of the phase 1 and 2 of the HMP (300 subjects). We found at least one sequencing read mapping to the SA_ASR sequence in 348 of the samples (27.5%) isolated from 110 (36.3%) of the subjects (Supplementary Figure 2). The presence of the species was variable across body sites, most commonly found in the left and right retroauricular creases and anterior nares (100%, 90% and 57%, respectively) and least common in the stool and subgingival plaque (6% and 5%, respectively) (Table 2). While the presence of the *S. aureus* reads in a sample will be dependent on factors such as the complexity of the microbiome and the amount of sequence data collected, this result was in line with estimates of *S. aureus* presence based on bacterial culture[15].

Of the 348 *S. aureus* positive samples, 321 had a *S. aureus* core coverage > 0.025X (Table 2). *S. aureus* was more prevalent at this level of coverage in the anterior nares, retroauricular creases and tongue dorsum. 165 (51%) of these samples were dominated by one subtype (largest ***β*** value > 0.8). In the other samples where there was a mixture of dominant subtypes, we used a conservation cutoff for a subtype being present as at least a minor component if the ***β*** value was >0.2 (chosen to conservatively remove overcalls due to random errors in sequence reads). Based on these definitions, the most commonly detected subtypes were CC30, CC8, CC45, CC398 and CC5 (present in 112 (35%), 72 (22%), 32 (10%), 29 (9%) and 26 (8%) samples, respectively) (Figure 4). There were only 2 subtypes not detected in any of the samples: CC123, and CC49. The US origin of the samples was reflected by the common occurrence of ST8 (representing lineages such as USA300 and USA500). CC30 and CC45 are common colonization isolates worldwide, whereas CC239 and CC22; prevalent in hospitals in Europe, Asia and Latin America [31][32][33,34], were infrequent in this study (present in 1 and 2 samples, respectively).

**Figure 4.**
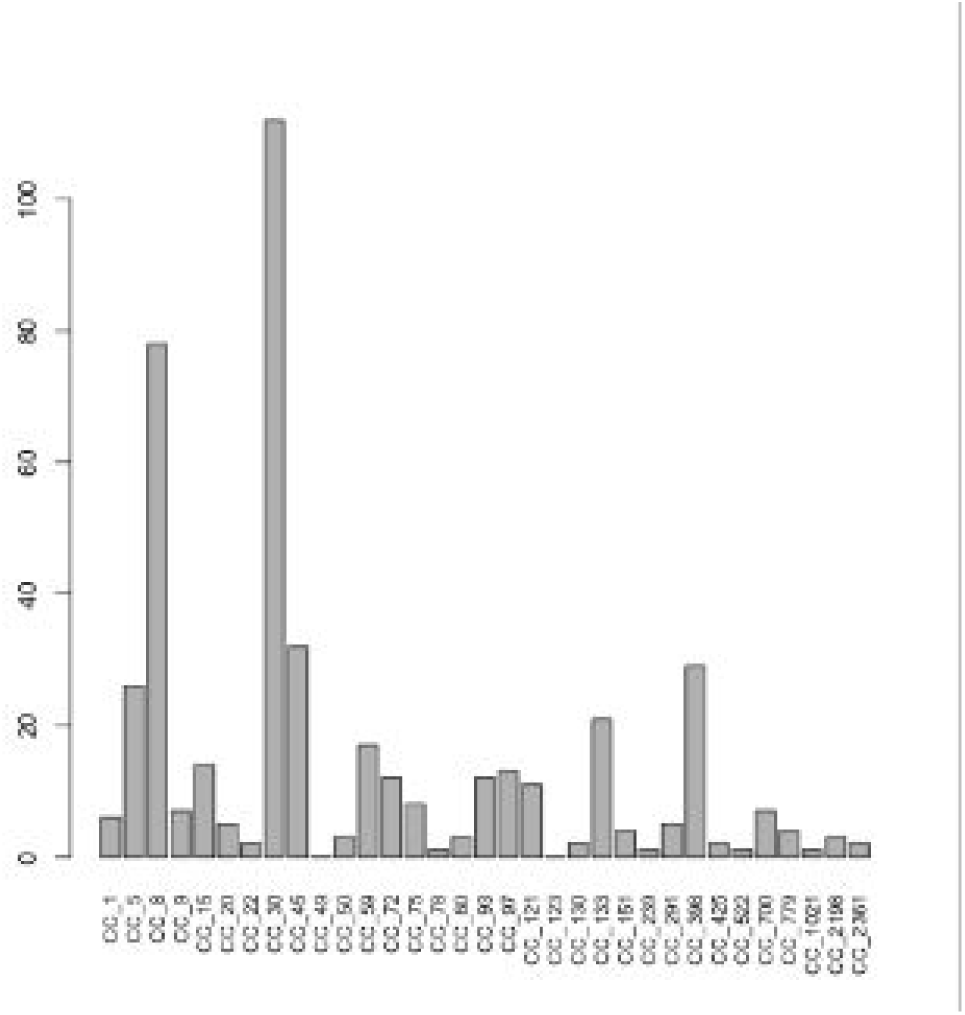
Distribution of subtypes (beta > 0 2) in S. aurous HMP samples with at least 0.025X genome coverage.

The median genome coverages of samples from the anterior nares, tongue dorsum and retroauricular creases were higher than that of the other body sites (with the exception of the hard palate where n=1; Supplementary Figure 3). However, in all but 21 out of 321 samples (93%) *S. aureus* genome coverage was below 2X. A sample from the anterior nares (NCBI SRA accession SRS011105) was a striking exception, with > 70x coverage of a CC59 S. *aureus* genome. Although the subtype schema was based on the core genome, we also calculated the coverage of the accessory *mecA* gene, necessary for MRSA. 55/1263 (4%) samples had reads mapping to *mecA.* There was no consistent relationship between the coverage level of the *S. aureus* genome and the coverage of *mecA,* even when we looked at the adjusted coverage (*binstrain* ***β*** multiplied by total coverage) for individual subtypes. This is likely because *S. aureus* subtypes can consist of both MRSA and non-MRSA strains (MSSA) [22,35] and *mecA* genes occur in *S. epidermidis* and other staphylococci [36] that may also be present in the sample. There were no *mecA* reads present in accession SRS011105, indicating that the CC59 strain present at 70x coverage was MSSA.

### *S. aureus* subtype distribution across subjects and body sites

We were interested to see if there were patterns to the distribution of *S. aureus* subtypes between different body sites in the HMP data set. Exploration of the incidence of common subtypes based on either the presence at a cutoff of ***β*** 0.2 or higher, or having the highest ***β*** value (Figures 5.a-c), appeared to show differences between body sites. For example, CC30 dominated the tongue dorsum, whereas CC8 was most prevalent in the buccal mucosa. The most striking bias in distribution was seen in the livestock associated CC398 subtype, which was found in the tongue dorsum in 26/29 (90%) samples (Figure 5c) where it was counted as present with a ***β***> 0.2. In order to understand the result, and as an example for learning about the type of evidence that produced positive subtype calls, we looked in more detail at the data underlying this call. We found that all the positive body sites samples contained, in addition a small number of rare SNPs, a synonymous SNP within the RNA polymerase gene that is conserved in almost all CC398 strains (Figure 6) but rarely found outside this subtype. This SNP was missing, or at low abundance from CC398 ***β*** < 0.2 samples.

**Figure 5.**
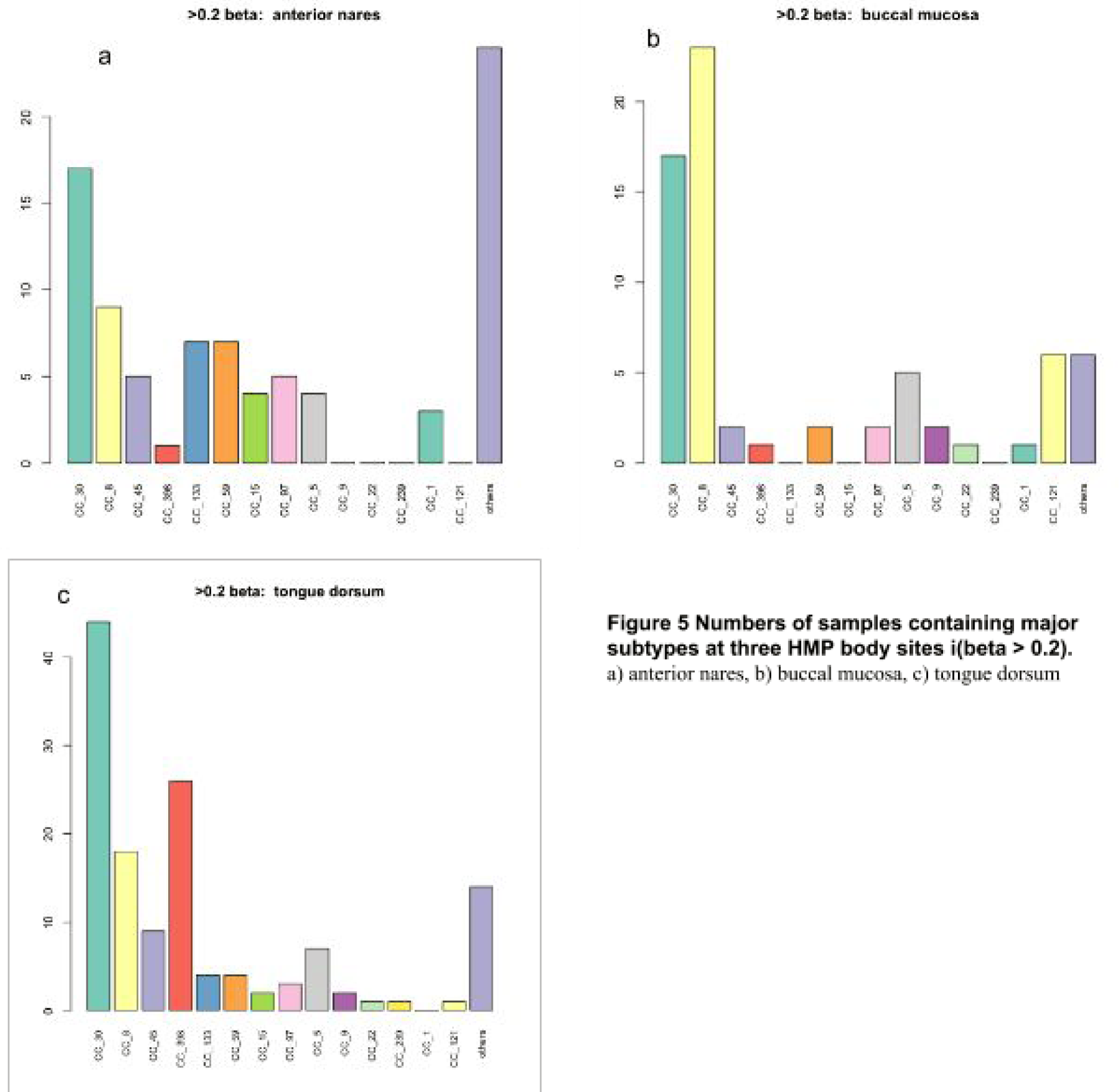
Numbers of samples containing major subtypes at three HMP body sites i(beta > 0,2). a) anterior nares, b) buccal mucosa. c) tongue dorsum

**Figure 6.**
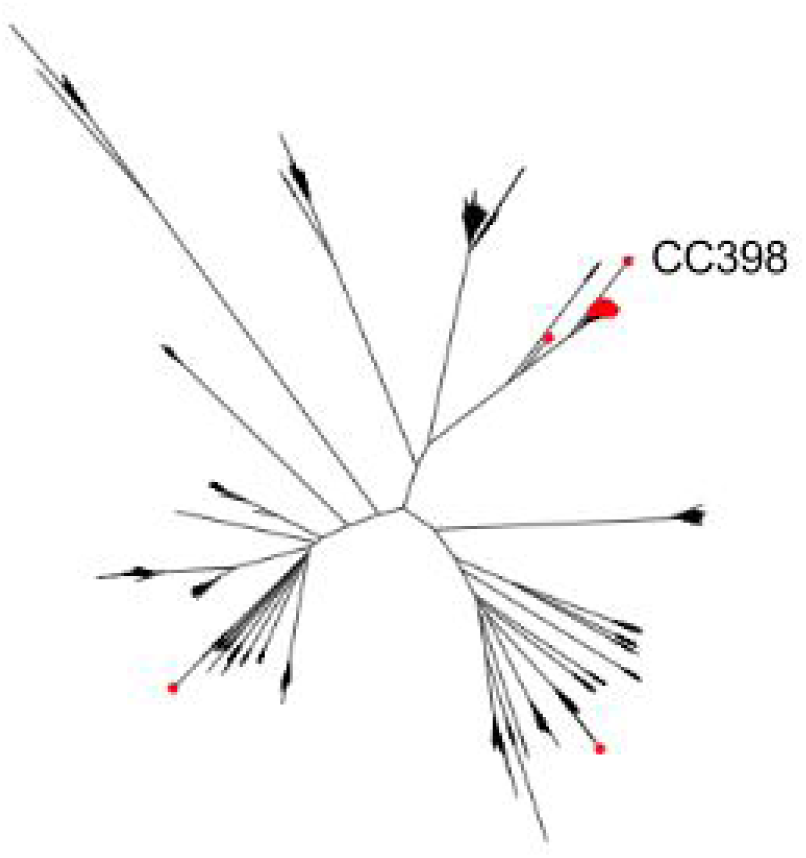
Phylogenetic placement of SNP responsible for calling CC398 in HMP samples. The synonymous SNP in the RNA polymerase gene is found primarily in strains of CC39B subtype. with the exception of two outlying strains.

Each sample was scored with a value of 1 for every subtype with a ***β*** value > 0.2 and then a distance matrix based on Hamming scores between samples was created. Using PERMANOVA, found there was a significant association between Hamming score distance and body site (p < 0.001, df = 8) but the model explained only ~11% of the variance. However, the result was tempered by finding a significant difference in beta dispersion between body sites (p < 0.001), which might have caused spurious association. We repeated the analysis using Hamming matrices based on having no minimum ***β*** value cutoffs and obtained similar results. As an alternative approach, we compared the sum of Hamming distances between samples of the same body type to 10,000 randomly selected samples (Table 3). The posterior fornix, buccal mucosa, stool and supragingival plaque had total Hamming scores that were < 5% of those found in random sampling.

**Table 3.** Within-site Hamming score compared to random permutations of all samples

| Body site | Number of samples(minimum 4) | p-value (compared to 10,000 permutations) |
| --- | --- | --- |
| Anterior nares | 67 | 0.50 |
| Buccal mucosa | 54 | 0.0008 |
| Left retroauricular crease | 23 | 0.85 |
| Palatine tonsils | 6 | 0.62 |
| Posterior fornix | 9 | 0.007 |
| Right retroauricular crease | 28 | 0.99 |
| Stool | 7 | 0.035 |
| Supragingival plaque | 37 | 0.0008 |
| Tongue dorsum | 82 | 0.78 |

Two body sites on the same individual may be more similar to each other than the same sites in other individuals because of intrapersonal spread of bacteria. There were 108 individuals within the HMP group with at least one body site containing > 0.025X *S. aureus* genome coverage. Using PERMANOVA based on Hamming distances and a subtype cutoff ***β*** value > 0.2 there was a significant association between subject and *S. aureus* composition (p = 0.005, df = 107, R^2^ = 0.4). Unlike the body site categories tested above, there were no significant differences in dispersion of Hamming distances between individuals. An interaction model for body site and subject accounted for ~47% of variance and had a p value of 0.001. However, when we instead based subtype composition without an arbitrary ***β*** value cutoff we lost significance in the interactive model (although the subject only p value was still < 0.05). As an alternative measure, we calculated the total Hamming distance of all *S. aureus* communities within the same individual (including both Phase I and Phase II data) and found it was lower than 10,000 random replicates of the same number of samples.

### Biotic and abiotic factors associated with *S. aureus* and its subtypes

The HMP collected abundant metadata on the participating subjects and we performed epidemiologic modeling using generalized linear mixed models to assess whether any variables were associated either with the presence of *S. aureus,* or with a specific subtype. In order to increase power we aggregated body sites into three categories: airways (anterior nares), oral cavity (attached keratinized gingiva, buccal mucosa, palatine tonsils, saliva, supragingival plaque and tongue dorsum) and skin (right and left retroauricular crease). In the binary outcome logistic regression model, main body site (p-value <0.001), having health insurance or not (p-value = 0.0525) and BMI (p-value = 0.0276) were significant predictors for the detection/presence of *S. aureus,* whereas for the multinomial logistic regression model with 4 outcomes, only main site (p-value = <0.001) and BMI (p-value = 0.0251) were significant predictors of the presence of *S. aureus* at significance level of 0.1 (Table 4). The estimated odds ratio for detecting the presence of any *S. aureus* subtypes in the airways compared to the oral cavity was 3.3 (95% CI: 2.2 - 5.0). This is consistent with our study and other previous studies showing that *S. aureus* is highly enriched in the anterior nares (nose) compared to any other body sites, and also body site could be a strong predictor for the presence of *S. aureus*.-In the multinomial model where specific subtypes were examined, odds of detection of CC8, CC30 and other subtypes were all significantly elevated in the airways compared to the oral cavity (Table 4). Similarly the odds of detecting any *S. aureus* subtypes in subjects with higher BMI was 70% higher when compared to subjects with normal BMI. In the multinomial model, odds of detection of CC8, CC30 and other subtypes were all elevated for higher vs. normal BMI, but the higher odds of detection was more pronounced for the other subtype group (OR CC8=1.4, 95% CI=0.6-3.0; OR CC30=1.1, 95%CI=0.5, 2.2; OR other subtype=2.4, 95% CI=1.3-4.5). Also the odds of detecting CC8 subtype tended to be higher in high BMI subjects while CC30 subtypes appeared to be associated with lower BMI. The binary outcome model suggested that the odds of detecting the presence of any *S. aureus* subtypes is 50% lower in those subjects that do not have health insurance than in those with health insurance. Even though race and ethnicity was not a statistical significant predictor for the detection of *S. aureus*, there was some indication that the odds of identifying any *S. aureus* subtype was higher among Hispanics compared to Non-Hispanic whites, with the odds ratio most elevated for detection of CC8 in the multinomial model (Table 4). However, we note that these odds ratios were based on only 45 samples from Hispanics included in our analysis. Our analysis also confirmed previous results that gender was not a significant predictor for the detection of any *S. aureus* subtypes. The odds of detecting *S. aureus* in females were 10% less than in males (Table 4). Age, breast fed or not, tobacco use (also previously identified in [17] and history of surgery were not significant predictors of association of the presence of *S. aureus* subtypes in any human body sites.

**Table 4.**
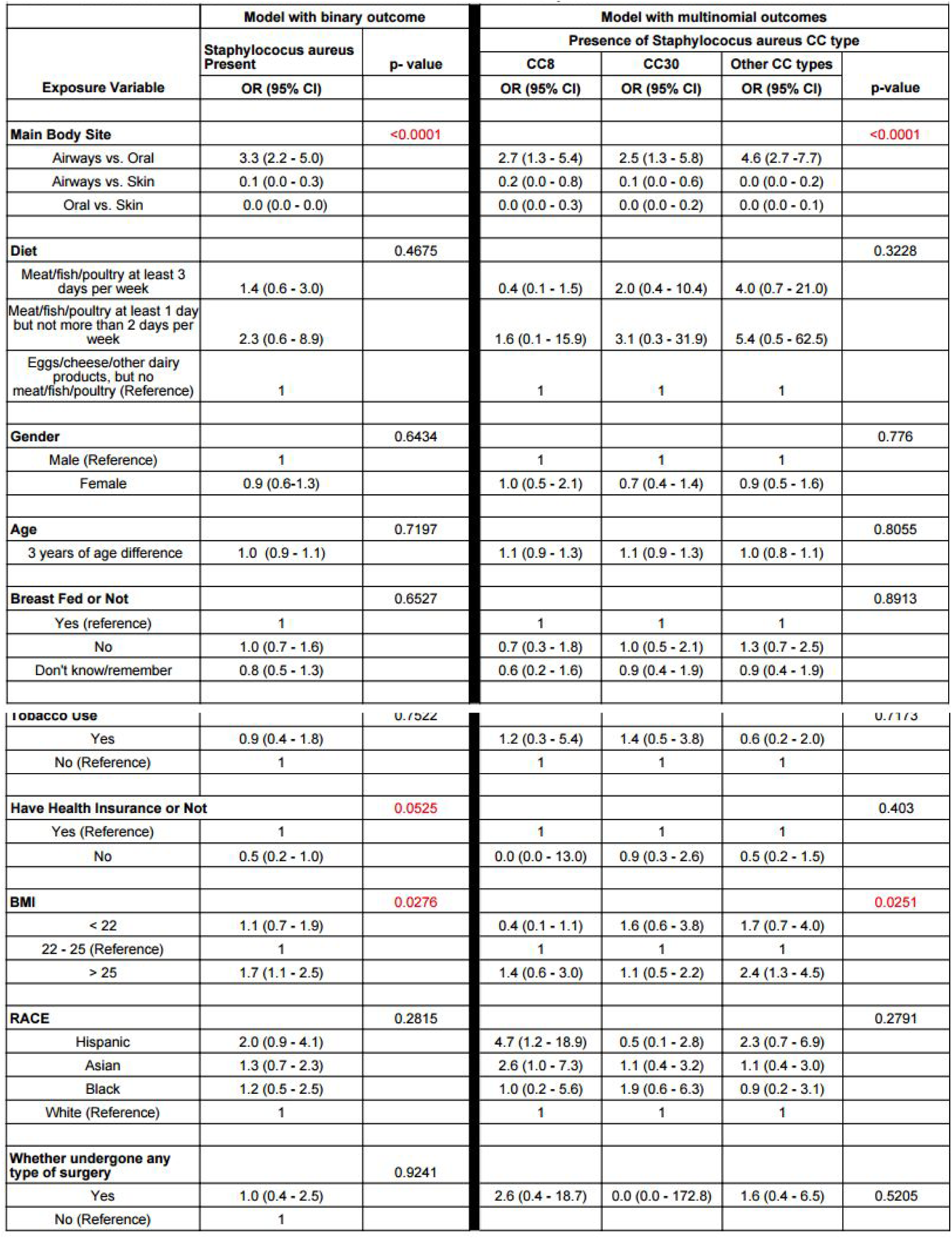
Estimated odds ratios with 95% confidence interval for models with binary outcome as well as multinomial outcomes

A major limitation of our epidemiologic analysis was the imprecision in our estimated odds ratios for detection of *S. aureus* driven mainly by the small number of study subjects and to some extent by the homogeneity of the HMP population for factors such as age, race, tobacco use, health insurance, diet, and history of surgery **(see Supplemental Table 3).** Despite this, we did observe evidence of more detection of *S. aureus* among subjects with higher BMI compared to normal BMI and suggestion of lower detection among subjects without health insurance.

### Geographical distribution of *S. aureus* subtypes found in the New York City subway setting

We looked at the geographic distribution of *S. aureus* subtypes in New York City (NYC), based on data provided in a recent environmental survey [37]. Afshinnekoo et al sampled 1,457 surfaces at all 466 open subway stations in NYC between mid 2013 to February 2014, while at the same time collecting detailed metadata using a customized iOS app. DNA extracted from the samples was shotgun sequenced using Illumina paired-end technology. We used our subtyping methodology to investigate *S. aureus* diversity within the study set, finding 149 of the metagenome samples to contain > 0.025X coverage of the *S. aureus* core genome. At a ***β*** value > 0.2 cutoff, CC30 and CC8 (29(20%) and 26(19%), respectively, were again the most common subtypes (Figure 7). CC5 (17(11%)) and CC45 (12(08%)) were also common, as we found in the HMP data, but CC2198 (6(4(10%)%)) and CC1 (9(6%)) were more prevalent than HMP.

**Figure 7.**
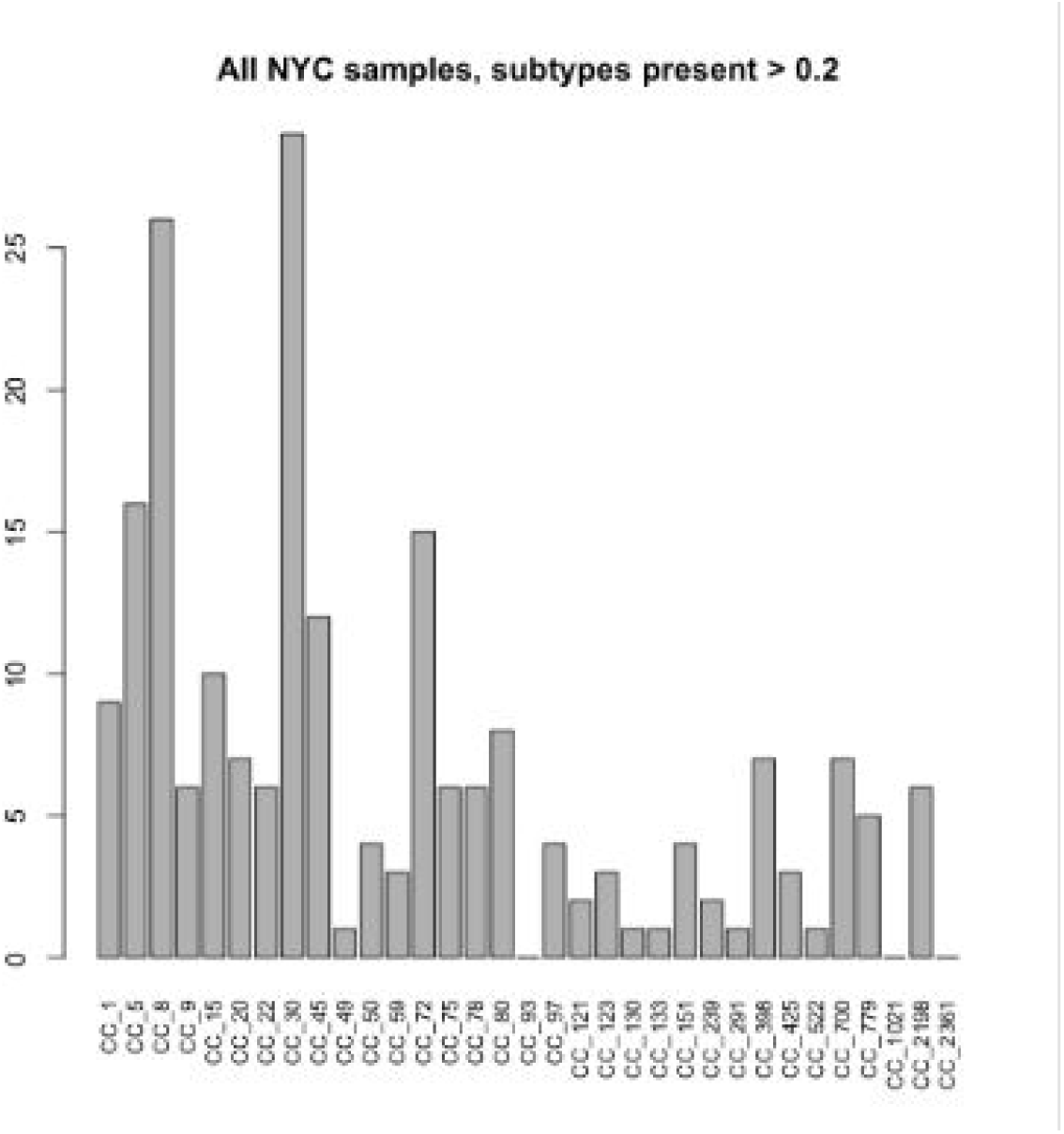
Numbers of samples from NYC subway with subtypes (beta > 0.2).

The NYC metadata allowed us to ask whether there was any geographical structure to the *S. aureus* population that might reflect compartmentalisation of the city based on ethnicity or other reasons. Afshinnekoo *et al* used the residual human DNA sequence in the metagenome data to show partitioning of areas of the city by race-specific polymorphisms that aligned with demographic data. We found little evidence of geographical clustering. Sites containing *S. aureus* were distributed more or less randomly over the New York City map, as were individual subtypes (Figure 8). Of the six most common subtypes, there was only one case (CC8) where the mean geographical distance between sites was in the lower 5% of NYC sites sampled at random (10,000 permutations). There was a significant regression between Hamming distance between sites and geographic distance (p < 0.001) but the effect size was small (R^2^< 1%). These result is in line with a recent study by Uhlemann et al [38], who found there was no strong geographic signal in the phylogeny of 387 strains of USA300 (CC8-ST8) isolated in households in Manhattan. These results suggest movement of *S. aureus* subtypes around the city may occur over a shorter timescale than the movement of the human population.

**Figure 8.**
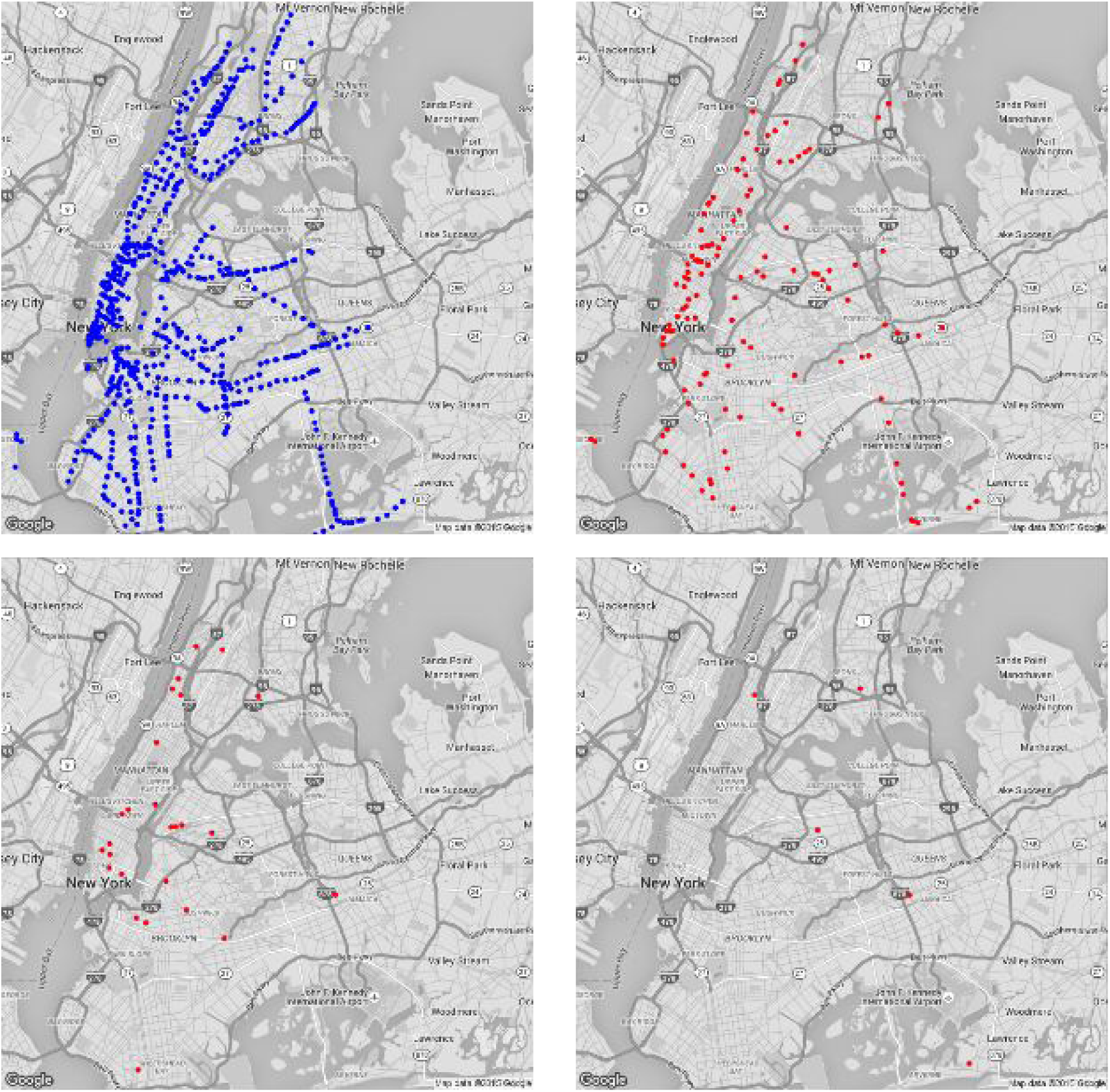
a) All sites sampled by Afshinnekoo et al 2015, b) sites containing *S. aureus* (beta > 0.2), c) sites containing CC30, d) sites containing CC22. Maps © Google.

## Discussion

In this work we developed a test to estimate the subtype profile of an important bacterial pathogen (*S. aureus*) from raw metagenomic data. We showed different mixtures of subtypes in different human subjects, and body sites and also gained an view of the the biogeography of *S. aureus* subtypes from environmental metagenomic data. The *binstrain* method should work well for studies seeking to examine variation in other bacterial species.

In the time period we worked on this project (2013-2015) there have a number of publications describing new software dedicated to taxonomic analysis of shotgun metagenomic data, reflecting the intense interest in this field [39,40], Several methods use unsupervised clustering of sequence reads belonging to the same species based on patterns of coverage covariance. Strategies include use of self-organizing maps[41], and co-abundance gene groups[42] [43][44][45]. These methods require processing multiple metagenomic sets simultaneously and are not explicitly designed for subtyping, although they could potentially be adapted for that purpose. The supervised strategies attempt to assign individual reads to species based on existing sequences stored within databases, using alignment based methods [46][47][48][49]. Kraken [50], CLARK[51] and OneCodex[52] use k-mer composition to efficiently place individual reads to their lowest common ancestor in the NCBI taxonomy hierarchy, kalisto is a fast, pseudo-alignment method, originally developed for RNA-seq analysis[53]. The phylosift pipeline [54] relies on read alignment to a reference genome followed by phylogenetic placement. A number of tools have been developed to type bacterial “strains” (ie below the species level). Snowball is an assembly-based method for strain resolution[55]. Latent strain analysis (LSA), a k-mer based approach, which does not require metagenome pre-assembly, is able to separate reads to the subspecies level[56] based on covariance. ShoRAH[57] attempts to estimate strain diversity based on mapping individual sequence reads, as does ConStrains[58]. WG-FAST [2] attempts phylogenetic placement based on assigning metagenomic reads from the species core genome that is effective for low coverage but hindered by strain mixtures. PathoScope[59] and Sigma[60] use SNP patterns from limited collections of references. This paragraph does not list every software in this fast-growing field and we refer readers to an online resource [40] to keep abreast of the latest developments in metagenome-based pathogen identification.

*binstrain* has a combination of features not contained in any other method. It explicitly models the presence of mixture of bacterial subtypes in the metagenome sample. The output is the assignment of metagenome data into bins that are convenient for microbiologists to interpret. Other programs, such as Kraken, can detect the signal of multiple strain of a species, but their output would need to be parsed by a post-processor in order to estimate the relative abundance of subspecies population groups. The distinction between assignment to individual reference “strains” and subtypes, which represent populations of strains, is important. The set of reference strains of a bacterial species in public databases is usually biased towards a small number of genotypes (often high-consequence pathogens). This can lead to an uneven picture of the actual composition of the species when using reference-based assignment. We were able to put together a less biased, population-based assignment of *S. aureus* by drawing on the unusually large number of diverse genomes sequences available in the public domain. However, even with more than 24,000 *S. aureus* genome projects in the public domain, there are still gaps in the coverage of rare genotypes that need to be covered. Unsupervised, metagenome strain discovery methods (e.g [42,56,58]) may play an important role in the future for validating how well subtyping methods capture the true extent of species population structure. The sensitivity of *binstrain* assigning subtypes (in terms of accurate assignment at low genome coverage) is set by the amount of nucleotide variability in the species and the number of subtypes. It can be expected that in many cases, as we showed for the two datasets here, there will be less than 2X genome coverage of the target species in the metagenome sample. The current version of the *binstrain* has some limitations. It examines individual SNPs independently and therefore does not use the information from SNPs co-occuring on the same sequencing read or on paired-end reads. The software does not take into account SNPs not already incorporated in the SNP matrix that may come from a novel subtype, and cannot detect possible recombination events (*S. aureus* is known to have a low rate of homologous recombination of < 1 kb pieces in its core genome[61] although occasional larger recombination events have been reported [62]). These issues will be addressed in the future iterations of the software, *binstrain* design considerations were developed for the data produced by the ubiquitous Illumina MiSeq and HiSeq instruments, which offer short reads (paired end sequences of up to 300 bp each) with low error rates [63], The future increase in metagenomic data sets produced using Pacific Biosciences and Oxford Nanopore technologies with long reads (typically > 5 kb) but higher error rates (typically 5-20%) will offer new opportunities (many more subtype specific SNP positions on the same sequence read) and challenges (obfuscation of the taxonomic signal due to errors).

Using the *binstrain* typing method we examined 321 samples from the HMP that contained *S. aureus* genome coverage > 0.025, the cutoff we established based on extensive testing of public *S. aureus* dataset. These were taken from 108 human subjects. Most culture-based studies of *S. aureus* carriage typically isolate a bacterial single colony from a small number of body sites (often just the nares). The study therefore represented an unusually large and extensive epidemiologic and genetic sampling of *S. aureus* carriage in healthy individuals.

We found evidence that the *S. aureus* subtype composition of different body sites in the same individual was more similar than would be expected from comparing random samples of the population. The caveat is that we only compared absolute presence and absence of subtypes instead of relative abundance and we used arbitrary **β** value thresholds. Further testing is necessary to refine the parameters, ideally by comparing the *binstrain* predictions to the results of culture-based studies. Nevertheless, the result is in line with the hypothesis that *S. aureus* spreads between body sites of the same individual [19], 49% of the samples contained mixtures of different subtypes, which would occur if there were multiple waves of colonizations followed by persistence.

The evidence for association of subtypes with different body sites was less conclusive. Despite the significant PERMANOVA p-value, the result was confounded by large differences in the variance of subtype composition between body sites. The permutation analysis (Table 3) showed that some body sites appeared to have more similar composition than expected by chance If a subtype was more associated with a human body this might be a the result of specific tissue-specific adaptation that leads to enhanced persistence following introduction. Body site specificity might also arise if different subtypes were unequally distributed in environments outside the body and thus were more likely to arrive on the human at different relative abundances depending on the route of entry, for example, whether by ingestion rather than hand contact. Despite the somewhat contradictory picture of body site specificity overall, one quite striking result was finding 90% of farm animal-associated CC_398 subtype [13,64] in the tongue dorsum site. It is tempting to speculate that this subtype was primarily acquired through ingestion of food products and it will be interesting to see if differences in *S. aureus* community composition between the tongue and other sites are found in other studies.

Of the epidemiologic variables available through the HMP, only body mass index (BMI) > 25 and possession of health insurance were associated with the presence of *S. aureus.* The subjects chosen for this study of the healthy human microbiome were fairly homogenous in terms of age, ethnicity [65], and absence of medical conditions, leaving little power to associate with conditions more prevalent in the general population. High BMI and health insurance as risks for *S. aureus* presence may be connected to healths factors outside those collected directly. There were no strong links to the carriage of the two major subtypes, CC30 and CC8, and there were no geographical distinction between the two in New York City subway station samples. Understanding the reasons behind the distributions of *S. aureus* subtypes will take larger data sets.

This study illustrated how well-annotated metagenomic datasets in the public domain are a rich data source that can be mined repeatedly for results beyond the original purposes for which they were created. *S. aureus* subtyping can be performed on any metagenomic dataset. Given the current growth in metagenomic sequence production, these could soon number in the hundreds of thousands to millions. Large-scale analysis subtype would give a much richer insight into *S. aureus* biogeography and colonization preferences than we have been able to achieve through conventional microbial sampling alone, a method which has been traditionally focused on one body site (anterior nares) and rarely takes into account the within sample genetic mixture of the species [66]. The *binstrain* method, and/or alternative approaches described above, can be extended to species other than *S. aureus*, giving us more tools to explore the bacterial diversity of this planet.

## Materials and Methods

### Classifying *S. aureus* subtypes based on a binomial mixture model

We used *binstrain* software [24], implemented in the R language [67] to perform *S. aureus* subtype classification. *binstrain* used a binomial mixture model to estimate the proportion of subtypes based on a DNA alignment against a reference (SA ASR, described below) and a matrix of SNPs that distinguish different genetic subtypes (construction of the matrix described below). *binstrain* assumes a binomial probability distribution, *p_i_* of observing a SNP, *x_i_* in the entire genome and *n_i_* denotes the total nucleotide coverage at position *i. Z_i,_*_j_ is an indicator function specifying whether *j^th^* strain has a SNP at *i’^h^* position. In the final version of the classifier, we used 102,057 SNP positions across the genome to classify *S. aureus* into 33 subtypes.

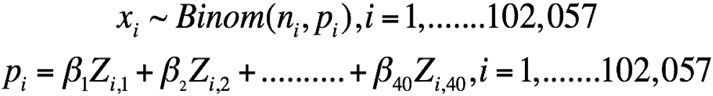

The estimation of **β***_i_* indicates the proportion of *S. aureus* reference strain-specific SNPs present in a clinical or purified sample. At the strain–specific SNP positions, there will be only a few ***β****_i_s* that affects *p_i_*. Other ***β****_i_ s* have no impact on *p_i_* because their corresponding *Z_ij_* are 0’s, which makes it a sparse design matrix. We utilized this sparsity of the design matrix in order to perform a well-established step-by-step procedure to estimate all the **βi** s using quadratic programming [24].

### *S. aureus* ancestral sequence regeneration

In order to elucidate the population structure of *S. aureus* and to select distinct subtypes, we performed ChromoPainter (http://www.paintmychromosomes.com) and fineSTRUCTURE [26] [68] analysis on 43 completed genomes downloaded from NCBI (Supplementary Table 1). The ChromoPainter analysis was applied using the linkage model to the genome-wide haplotype data generated from a whole genome progressive MAUVE [69] alignment of the 43 genomes to generate a coancestry matrix, which was used by the fineSTRUCTURE algorithm to perform model-based clustering using the Bayesian MCMC approach. This analysis indicated the presence of 19 distinct populations of *S. aureus* out of the 43 completed genomes used (Supplementary Table 1). We selected one genome sequence from each of the 19 groups (choosing randomly in the cases where more than one genome sequence was represented) and reconstructed the phylogenetic tree using the maximum likelihood approach by RAxML [70] with 100 bootstraps based on our MAUVE alignment. The ancestral sequence of *S. aureus* (SA_ASR) (2,872,915 bp) (Supplementary Data 1) was generated using the baseml program in the PAML package [71] with the whole genome alignment and phylogenetic tree as inputs. We implemented GTR nucleotide substitution model, with 5 gamma rate categories, assuming that the model was homogeneous across all the sites.

### Development of a *S. aureus binstrain* SNP matrix

We initially constructed a *binstrain* SNP matrix [24] based on the 19 reference genomes of the SA_ASR. We tested this prototype classifier against 2,692 diverse *S. aureus* genomes downloaded as FASTQ files from NCBI (Supplementary Data 2). We first mapped the reads from each genome against the SA_ASR to generate the base call and coverage (average read depth) in each position in the mpileup output format. The alignment was performed using the Burrows-Wheeler Aligner (BWA) (Version: 0.6.1-r104) short-read aligner[72] by specifying the maximum number of gap extensions (e) to be 10. The resultant short-read alignment files for each samples were converted to mpileup format using the mpileup option in SamTools software along with the –B option that disables probabilistic realignment for the computation of base alignment quality (BAQ). When challenged using *binstrain* we found that only 1379 of the 2692 (51.22%) genomes estimated a **β** value >=0.8 indicating that our sample of *S. aureus* genomes did not represent the diversity of the species. We constructed a second version of the SNP pattern file based on a whole genome progressive MAUVE alignment with an additional 21 draft *S. aureus* genome chosen for diversity (Supplemental Table 2). We filtered out all the SNP positions where any of the *S. aureus* strains and two genomes of the near-neighbor *S. epidermidis* species[73] *(S. epidermidis ATCC12228* (accession number: NC_004461.1) and *S. epidermidis RP62A* (accession number: NC_002976.3) had a SNP at the same position. The resultant final SNP pattern file contained a total of 102,057 SNP positions across the entire core genome. The SNP pattern file contained SNPs unique for each reference strain as well as SNPs shared among more than one reference strains represented in this study. To determine specificity and sensitivity of the new classifier we tested the 2114 diverse *S. aureus* strains from the NCBI SRA database with ST determined by SRST software[28], downsampled to genome coverage between 0.025X and 0.75X (Supplemental data 2).

### Testing subtype classification using simulated sequence data

In order to test the *S. aureus* subtype classification scheme we used ART software[29] to simulate single end 100 bp Illumina read sequence data based on assembled *S. aureus* genome templates. ART simulates sequencing reads by mimicking the output of the Illumina sequencing process with empirical error models summarized from large recalibrated sequencing data. FASTQ files were generated based on the 40 *S. aureus* reference genomes used in this study at varying coverages. A total of 3600 FASTQ files were simulated for each strain 9 times separately at the following genome coverages (X): 0.000625X, 0.0125X, 0.025X, 0.05X, 0.1X, 0.5X, 0.75X, 1X, 2.5X and 5X. In addition, we simulated 3 artificial microbiome samples by individually simulating sequencing reads of the major bacteria present in a healthy anterior nares identified by the HMP [30] and concatenated them all together with 3 random *S. aureus* reference genomes (CC_151, CC_239 and MLST_93) (Figure 1). The non *S. aureus* bacteria simulated were *Propionibacterium acnes* (200X), S. epidermidis (100X), *Streptococcus mitis* (7X), *Bacteroides vulgatus* (5X), *Staphylococcus haemolyticus* (4X) and *Staphylococcus saprophyticus* (3X). Following are the proportions of the 3 *S. aureus* genomes in each of the 3 simulated microbiome samples: **1) microbiome 1** – CC_151_151 (0.5X), CC_239_239 (0.1X) and MLST_93 (0.25X) **2) microbiome 2 –** CC_151_151 (1.5X), CC_239_239 (1X) and MLST_93 (0.75X) and 3) **microbiome 3** – CC_151_151 (1.0X), CC_239_239 (5.0X) and MLST_93 (2.5X).

### Sequence data analysis and statistical modeling for SNP based genotyping of *S. aureus* strains in the HMP and NYC subway metagenome projects

For this study we obtained raw mwgs sequence data in fastq files for a total of 1265 samples (human DNA removed using NCBFs BMTagger tool) from the HMP ftp site (ftp://public-ftp.hmpdacc.org/Illumina). The HMP carried out 2 phases of metagenomic whole genome shotgun sequencing (mwgs), performed using the Illumina GAIIx platform with 101 bp paired-end reads. For Phase 1, 764 samples were chosen from 103 adults and for Phase II, 400 samples were chosen from 67 adults. Samples were chosen covering 16 body sites. The Phase 1 data sets have been described previously [30] [74], We also used mwgs data from DNA samples extracted from the New York subway [37], The 125 x 125 bp paired end sequence reads were prepared using an Illumina HiSeq 2500 instrument. Sequence data was downloaded from the NCBI SRA repository (project PRJNA271013).

We mapped the FASTQ files from these projects at against the SA_ASR to obtain mpileup format data using BWA as described above and called the subtypes present using *binstrain.* From the mpileup output we calculated the average gene coverage mapping to the *S. aureus* core genome. In addition to the core genome, we also determined the number of reads mapping to a representative *S. aureus mecA* gene (Locus AKR51832.1 gi:899756207).

### Selection of metadata from the HMP and epidemiological modelling

A large amount of demographic and clinical data were collected for each of the individuals sampled for the HMP. We obtained access to the most recent version of these data through a formal request to the dbGap database (accession phs000228.v3.p1). We used the metadata from the 170 healthy individuals (phase I and II) from whom the HMP mwgs sequence data were generated. Because of the generally high level of health of the subjects, for most clinical variables there were too few cases to have realistic odds of association. The binary categorical variables (exposure variables) from the metadata, which we investigated in relation to presence of *S. aureus* and/or a particular *S. aureus* subtype in a body site were gender, breastfed or not, tobacco use, insurance information and history of previous surgery (Supplementary Table 3). Other categorical variables used were diet (Meat/fish/poultry at least three days per week, Meat/fish/poultry at least one day but not more than two days per week and Eggs/cheese/other dairy products, but no meat/fish/poultry), race/ethnicity (Hispanic, asian, non-Hispanic black and non-Hispanic white), BMI (< 22, 22 - 25 & > 25). Age was treated as a continuous variable that ranged from 18 to 40 years of age (Supplementary Table 3). For the epidemiological analysis, we included only those body sites where *S. aureus* was detected in at least 20% of the samples within each of the body sites sampled for HMP, and also regrouped them into main body sites based on their proximity within the human body. The 3 main sites were; airways (anterior nares), oral cavity (attached keratinized gingiva, buccal mucosa, palatine tonsils, saliva, supragingival plaque and tongue dorsum) and skin (right and left retroauricular crease). There were a total of 840 samples collected from 133 participants in the HMP, used in the epidemiological analysis (described below).

We performed both binary and multinomial (with 4 outcomes) logistic regression to identify predictors for *S. aureus* detection among HMP participants. The binary outcome indicated whether the presence of *S. aureus* was detected or not detected (reference), while the 4 outcomes for the multinomial logit model were the presence/detection of *S. aureus* CC8, CC30, any other *S. aureus* CC types and no detection of *S. aureus* (reference). Odds ratios were estimated by fitting generalized linear mixed models using SAS PROC GLIMMIX (Version 9.4, Cary, NC) with main site and other exposure variables (described above) as fixed effects and random effects for subject in order to assess any possible association of the exposure variables and the presence/detection of *S. aureus.*

### Association of *S. aureus* HMP subtypes with body site and individual subjects

For each HMP project that contained *S. aureus* reads we prepared a Hamming distance matrix between samples based on the shared presence of subtypes inferred from the binstrain beta estimates. We only used subtypes with at least a beta value of 0.2, in order to mitigate the possible effect of overcalling subtypes. From this distance matrix we performed a principal components analysis and mapped the distribution of samples by body site and individual subject. In order to test whether individual body sites were more likely to contain similar subtypes we calculated the total Hamming distance between all samples of the same body type and compared the value to 10,000 random draws of the same number of samples from the total set of *S. aureus* positive samples.

### Co-occurrence of rare SNPs between body sites in an individual

We made a matrix of all variant positions in mpileup files generated for comparison of each metagenome to the SA_ASR described above. Initially we counted a variant supported by as few as one read. We then counted the incidence of each SNP variant across the 348 *S. aureus* positive samples on filtered out all but the variants present in 2-5 samples. (The upper limit of five was chosen as the maximum diversity of Staph-positive body sites present in one individual. We then created a co-occurrence matrix of variants shared samples and normalized to a binary, indicating whether or any two samples shared at least one rare SNP. We then counted the number of times pairs of body sites (e.g anterior nares/ buccal mucosa) in the same individual shared rare SNPs and used a Fisher’s exact test to determine if the number was enriched compared to the population of pairs of body sites found in different individuals.

### Spatial relationships in NYC subway sampling sites

We used a permutation test to ascertain whether a matrix of geographical distances between Staph-positive NYC sampling sites correlated with the matrix of Hamming distances of subtype *binstrain* beta values.

The analyses described in this paper made use of the R [67] packages, *binstrain [24], vegan, e1071, gdata, Rgooglemaps, dplyr* and *reshape2* and SAS PROC GLIMMIX (Version 9.4, Cary, NC).

## Supplemental data

<u>Supplementary Table 1:</u> The list of the 43 complete genomes of Staphylococcus aureus used for the fineStructure analysis. (http://dx.doi.org/10.6084/m9.figshare.1591908)

<u>Supplementary Table 2:</u> List of *Staphylococcus aureus* genomes used for the version 1(19 genomes) and version 2 (an additional 21 genomes) for constructing the SNP pattern file used in this study. (http://dx.doi.org/10.6084/m9.figshare.1591910)

<u>Supplementary Table 3:</u> List of all the exposure variables assessed in this study, which includes the demographics and characteristics of the participants. (http://dx.doi.org/10.6084/m9.figshare.1591919)

<u>Supplementary Table 4:</u> Specificity of the v2 subtyping matrix on 2114 *S. aureus* FASTQ files from the NCBI SRA. (http://dx.doi.org/10.6084/m9.figshare.1593023)

<u>Supplementary Figure 1:</u> Figure showing the binstrain estimates for the analysis on 10 simulated sequence read sets of between 0.00625X and 5 X genome coverage for each of the 40 Staphylococcus aureus subtype-defining genomes and repeated the analysis 9 times (a total of 3600 fastq files and binstrain runs). (http://dx.doi.org/10.6084/m9.figshare.1591909)

<u>Supplementary Figure 2:</u> Heatmap showing the binstrain beta estimates of the *Staphylococcus aureus* positive HMP samples. (http://dx.doi.org/10.6084/m9.figshare.1591909)

<u>Supplementary Figure 3:</u> Staphylococcus aureus genome coverage for the HMP samples at the various body sites.(http://dx.doi.org/10.6084/m9.figshare.1592157)

<u>Supplementary Data 1:</u> Estimated ancestral sequence of *Staphylococcus aureus* used for reference mapping in this study.(http://dx.doi.org/10.6084/m9.figshare.1591912)

<u>Supplementary Data 2:</u> Information of the 2,114 *Staphylococcus aureus* fastq files obtained from the NCBI SRA database to estimate the specificity and sensitivity of binstrain software, (http://dx.doi.org/10.6084/m9.figshare.1591918)

More details of the analysis can be obtained from the public github site: https://github.com/Read-Lab-Confederation/staph_metagenome_subtypes.

## Acknowledgements

This worked was funded through development funds from Emory University School of Medicine. Part of this work will be presented as a Master’s thesis at Rollins School of Public Health, Emory University by SJJ towards his MPH degree in Applied Epidemiology. Access to de-identified data was approved under dbGAP agreement #18287-4. “Funding support for the development of NIH Human Microbiome Project - Core Microbiome Sampling Protocol A (HMP-A) was provided by the NIH Roadmap for Medical Research.Clinical data from this study were jointly produced by the Baylor College of Medicine and the Washington University School of Medicine. Sequencing data were produced by the Baylor College of Medicine Human Genome Sequencing Center, The Broad Institute, the Genome Center at Washington University, and the J. Craig Venter Institute. These data were submitted by the EMMES Corporation, which serves as the clinical data collection site for the HMP. “

## References

1. Joseph SJ, Read TD. Bacterial population genomics and infectious disease diagnostics. Trends Biotechnol. 2010;28: 611–618.

2. Sahl JW, Schupp JM, Rasko DA, Colman RE, Foster JT, Keim P. Phylogenetically typing bacterial strains from partial SNP genotypes observed from direct sequencing of clinical specimen metagenomic data. Genome Med. 2015;7: 52.

3. King MD, Humphrey BJ, Wang YF, Kourbatova EV, Ray SM, Blumberg HM. Emergence of community-acquired methicillin-resistant Staphylococcus aureus USA 300 clone as the predominant cause of skin and soft-tissue infections. Ann Intern Med. Am Coll Physicians; 2006; 144: 309–317.

4. Seybold U, Kourbatova EV, Johnson JG, Halvosa SJ, Wang YF, King MD. SM Ray, and HM Blumberg. 2006. Emergence of community-associated 17 methicillin-resistant Staphylococcus aureus USA300 genotype as a major cause of 18 health care-associated blood stream infections. Clin Infect Dis. 16AD;42: 647–656.

5. Mera RM, Suaya JA, Amrine-Madsen H, Hogea CS, Miller LA, Lu EP, et al. Increasing Role of Staphylococcus aureus and Community-Acquired Methicillin-Resistant Staphylococcus aureus Infections in the United States: A 10-Year Trend of Replacement and Expansion. Microb Drug Resist. 2011; 17: 321–328.

6. O’Hara FP, Amrine-Madsen H, Mera RM, Brown ML, Close NM, Suaya JA, et al. Molecular Characterization of Staphylococcus aureus in the United States 2004-2008 Reveals the Rapid Expansion of USA300 Among Inpatients and Outpatients. Microb Drug Resist. 2012;18: 555–561.

7. McBryde ES, Bradley LC, Whitby M, McElwain DLS. An investigation of contact transmission of methicillin-resistant Staphylococcus aureus. J Hosp Infect. 2004;58: 104–108.

8. PhD JAO, PhD SY, French GL, FRCPath. The Role Played by Contaminated Surfaces in the Transmission of Nosocomial Pathogens •. Infect Control Hosp Epidemiol. The University of Chicago Press on behalf of The Society for Healthcare Epidemiology of America; 2011;32: 687–699.

9. Dulon M, Haamann F, Peters C, Schablon A, Nienhaus A. MRSA prevalence in European healthcare settings: areview. BMC Infect Dis. 2011; 11: 138.

10. Klevens RM, Morrison MA, Nadle J, Petit S, Gershman K, Ray S, et al. Invasive methicillin-resistant Staphylococcus aureus infections in the United States. JAMA. 2007;298: 1763–1771.

11. Kuehnert MJ, Kruszon-Moran D, Hill HA, McQuillan G, McAllister SK, Fosheim G, et al. Prevalence of Staphylococcus aureus Nasal Colonization in the United States, 2001-2002. J Infect Dis. 2006;193: 172–179.

12. Rinsky JL, Maya N, Steve W, Devon H, Dothula B, Price LB, et al. Livestock-Associated Methicillin and Multidrug Resistant Staphylococcus aureus Is Present among Industrial, Not Antibiotic-Free Livestock Operation Workers in North Carolina. PLoS One. 2013;8: e67641.

13. Price LB, Stegger M, Hasman H, Aziz M, Larsen J, Andersen PS, et al. Staphylococcus aureus CC398: host adaptation and emergence of methicillin resistance in livestock. MBio. 2012;3. doi: 10.1128/mBio.00305-11

14. Paterson GK, Harrison EM, Murray GGR, Welch JJ, Warland JH, Holden MTG, et al. Capturing the cloud of diversity reveals complexity and heterogeneity of MRS A carriage, infection and transmission. Nat Commun. 2015;6: 6560.

15. Van Belkum A, Verkaik NJ, de Vogel CP, Boelens HA, Verveer J, Nouwen JL, et al. Reclassification of Staphylococcus aureus nasal carriage types. J Infect Dis. 2009; 199: 1820–1826.

16. Lamers RP, Stinnett JW, Muthukrishnan G, Parkinson CL, Cole AM. Evolutionary analyses of Staphylococcus aureus identify genetic relationships between nasal carriage and clinical isolates. PLoS One. 2011;6: e16426.

17. Liu CM, Price LB, Hungate BA, Abraham AG, Larsen LA, Christensen K, et al. Staphylococcus aureus and the ecology of the nasal microbiome. Science Advances. American Association for the Advancement of Science; 2015; 1: e1400216.

18. Eiff C von, Becker K, Machka K, Stammer H, Peters G. Nasal carriage as a source of Staphylococcus aureus bacteremia. Study Group. N Engl J Med. 2001;344: 11–16.

19. Kluytmans J, van Belkum A, Verbrugh H. Nasal carriage of Staphylococcus aureus: epidemiology, underlying mechanisms, and associated risks. Clin Microbiol Rev. 1997; 10: 505–520.

20. Brown AF, Leech JM, Rogers TR, McLoughlin RM. Staphylococcus aureus Colonization: Modulation of Host Immune Response and Impact on Human Vaccine Design. Front Immunol. 2014;4: 507.

21. Feil EJ, Li BC, Aanensen DM, Hanage WP, Spratt BG. eBURST: inferring patterns of evolutionary descent among clusters of related bacterial genotypes from multilocus sequence typing data. J Bacteriol. 2004; 186: 1518–1530.

22. Nübel U, Roumagnac P, Feldkamp M, Song J-H, Ko KS, Huang Y-C, et al. Frequent emergence and limited geographic dispersal of methicillin-resistant Staphylococcus aureus. Proc Natl Acad Sci USA. 2008;105: 14130–14135.

23. Kobayashi SD, Musser JM, DeLeo FR. Genomic analysis of the emergence of vancomycin-resistant Staphylococcus aureus. MBio. 2012;3. doi: 10.1128/mBio.00170-12

24. Joseph SJ, Li B, Ghonasgi T, Haase CP, Qin ZS, Dean D, et al. Direct Amplification, Sequencing and Profiling of Chlamydia trachomatis Strains in Single and Mixed Infection Clinical Samples. PLoS One. 2014;9: e99290.

25. Darling ACE, Mau B, Blattner FR, Perna NT. Mauve: multiple alignment of conserved genomic sequence with rearrangements. Genome Res. 2004;14: 1394–1403.

26. Yahara K, Furuta Y, Oshima K, Yoshida M, Azuma T, Hattori M, et al. Chromosome painting in silico in a bacterial species reveals fine population structure. Mol Biol Evol. 2013;30: 1454–1464.

27. Feil EJ, Cooper JE, Grundmann H, Robinson DA, Enright MC, Berendt T, et al. How clonal is Staphylococcus aureus? J Bacteriol. 2003;185: 3307–3316.

28. Inouye M, Conway TC, Zobel J, Holt KE. Short read sequence typing (SRST): multi-locus sequence types from short reads. BMC Genomics. BioMed Central Ltd; 2012; 13: 338.

29. Huang W, Li L, Myers JR, Marth GT. ART: a next-generation sequencing read simulator. Bioinformatics. 2012;28: 593–594.

30. Human Microbiome Project Consortium. Structure, function and diversity of the healthy human microbiome. Nature. 2012;486: 207–214.

31. Baines SL, Holt KE, Schultz MB, Seemann T, Howden BO, Jensen SO, et al. Convergent Adaptation in the Dominant Global Hospital Clone ST239 of Methicillin-Resistant Staphylococcus aureus. MBio. 2015;6. doi: 10.1128/mBio.00080-15

32. Harris SR, Feil EJ, Holden MTG, Quail MA, Nickerson EK, Chantratita N, et al. Evolution of MRSA During Hospital Transmission and Intercontinental Spread. Science. 2010;327: 469–474.

33. Stanczak-Mrozek KI, Manne A, Knight GM, Gould K, Witney AA, Lindsay JA. Within-host diversity of MRSA antimicrobial resistances. J Antimicrob Chemother. 2015; doi:10.1093/jac/dkv119

34. Knight GM, Budd EL, Whitney L, Thornley A, Ghusein H Al-, Planche T, et al. Shift in dominant hospital-associated methicillin-resistant Staphylococcus aureus (HA-MRSA) clones overtime. J Antimicrob Chemother. 2012;67: 928–932.

35. Enright MC, Robinson DA, Randle G, Feil EJ, Grundmann H, Spratt BG. The evolutionary history of methicillin-resistant Staphylococcus aureus (MRSA). Proc Natl Acad Sci USA. 2002;99: 7687–7692.

36. Diekema DJ, Pfaller MA, Schmitz FJ, Smayevsky J, Bell J, Jones RN, et al. Survey of infections due to Staphylococcus species: frequency of occurrence and antimicrobial susceptibility of isolates collected in the United States, Canada, Latin America, Europe, and the Western Pacific region for the SENTRY Antimicrobial Surveillance Program, 1997-1999. Clin Infect Dis. 2001;32 Suppl 2: S114–32.

37. Afshinnekoo E, Meydan C, Chowdhury S, Jaroudi D, Boyer C, Bernstein N, et al. Geospatial Resolution of Human and Bacterial Diversity with City-Scale Metagenomics. Cell Systems. Elsevier; 2015; 1: 72–87.

38. Uhlemann A-C, Dordel J, Knox JR, Raven KE, Parkhill J, Holden MTG, et al. Molecular tracing of the emergence, diversification, and transmission of S. aureus sequence type 8 in a New York community. Proc Natl Acad Sci USA. 2014; doi: 10.1073/pnas. 1401006111

39. Lindgreen S, Adair KL, Gardner P. An evaluation of the accuracy and speed of metagenome analysis tools. bioRxiv. 2015. p. 017830.

40. Jacobs J. Pipelines for Pathogen Identification. In: Google Docs [Internet], [cited 2 Nov 2015], Available: https://docs.google.com/document/d/1qLczhk4MAKjkOtz-PnXhmgGEWnWQJHLfOMjd1B9U2RU/edit

41. Sharon I, Morowitz MJ, Thomas BC, Costello EK, Relman DA, Banfield JF. Time series community genomics analysis reveals rapid shifts in bacterial species, strains, and phage during infant gut colonization. Genome Res. 2013;23: 111–120.

42. Nielsen HB, Almeida M, Juncker AS, Rasmussen S, Li J, Sunagawa S, et al. Identification and assembly of genomes and genetic elements in complex metagenomic samples without using reference genomes. Nat Biotechnol. nature.com; 2014;32: 822–828.

43. Kang DD, Froula J, Egan R, Wang Z. MetaBAT, an efficient tool for accurately reconstructing single genomes from complex microbial communities. PeerJ. PeerJ Inc.; 2015;3: e1165.

44. Imelfort M, Parks D, Woodcroft BJ, Dennis P, Hugenholtz P, Tyson GW. GroopM: an automated tool for the recovery of population genomes from related metagenomes. PeerJ. PeerJ Inc.; 2014;2: e603.

45. Alneberg J, Bjarnason BS, de Bruijn I, Schirmer M, Quick J, Ijaz UZ, et al. Binning metagenomic contigs by coverage and composition. Nat Methods. 2014; 11: 1144–1146.

46. Naccache SN, Federman S, Veeraraghavan N, Zaharia M, Lee D, Samayoa E, et al. A cloud-compatible bioinformatics pipeline for ultrarapid pathogen identification from next-generation sequencing of clinical samples. Genome Res. 2014;24: 1180–1192.

47. Segata N, Waldron L, Ballarini A, Narasimhan V, Jousson O, Huttenhower C. Metagenomic microbial community profiling using unique clade-specific marker genes. Nat Methods. 2012;9: 811–814.

48. Freitas TAK, Li P-E, Scholz MB, Chain PSG. Accurate read-based metagenome characterization using a hierarchical suite of unique signatures. Nucleic Acids Res. 2015; doi: 10.1093/nar/gkv 180

49. Huson DH, Auch AF, Qi J, Schuster SC. MEGAN analysis of metagenomic data. Genome Res. 2007; 17: 377–386.

50. Wood DE, Salzberg SL. Kraken: ultrafast metagenomic sequence classification using exact alignments. Genome Biol. 2014;15: R46.

51. Ounit R, Wanamaker S, Close TJ, Lonardi S. CLARK: fast and accurate classification of metagenomic and genomic sequences using discriminative k-mers. BMC Genomics. 2015; 16: 236.

52. Minot SS, Krumm N, Greenfield NB. One Codex: A Sensitive and Accurate Data Platform for Genomic Microbial Identification. bioRxiv. 2015. p. 027607.

53. Schaeffer L, Pimentel H, Bray N, Melsted P, Pachter L. Pseudoalignment for metagenomic read assignment. arXiv [q-bio.QM], 2015.

54. Darling AE, Jospin G, Lowe E, Matsen FA 4th, Bik HM, Eisen JA. PhyloSift: phylogenetic analysis of genomes and metagenomes. PeerJ. 2014;2: e243.

55. Gregor I, Schönhuth A, McHardy AC. Snowball: Strain aware gene assembly of Metagenomes. arXiv [q-bio.QM], 2015.

56. Cleary B, Brito IL, Huang K, Gevers D, Shea T, Young S, et al. Detection of low-abundance bacterial strains in metagenomic datasets by eigengenome partitioning. Nat Biotechnol. Nature Publishing Group; 2015; doi:10.1038/nbt.3329

57. Zagordi O, Bhattacharya A, Eriksson N, Beerenwinkel N. ShoRAH: estimating the genetic diversity of a mixed sample from next-generation sequencing data. BMC Bioinformatics. 2011; 12: 119.

58. Luo C, Knight R, Siljander H, Knip M, Xavier RJ, Gevers D. ConStrains identifies microbial strains in metagenomic datasets. Nat Biotechnol. 2015;33: 1045–1052.

59. Hong C, Manimaran S, Shen Y, Perez-Rogers JF, Byrd AL, Castro-Nallar E, et al. PathoScope 2.0: a complete computational framework for strain identification in environmental or clinical sequencing samples. Microbiome. 2014;2: 33.

60. Ahn T-H, Chai J, Pan C. Sigma: strain-level inference of genomes from metagenomic analysis for biosurveillance. Bioinformatics. 2015;31: 170–177.

61. Castillo-Ramírez S, Corander J, Marttinen P, Aldeljawi M, Hanage WP, Westh H, et al. Phylogeographic variation in recombination rates within a global clone of Methicillin-Resistant Staphylococcus aureus (MRSA). Genome Biol. 2012; 13: R126.

62. Spoor LE, Richardson E, Richards AC, Wilson GJ, Mendonca C, Gupta RK, et al. Recombination-mediated remodelling of host-pathogen interactions during Staphylococcus aureus niche adaptation. Microbial Genomics. Microbiology Society; 2015; doi:10.1099/mgen.0.000036

63. Rosen MJ, Davison M, Bhaya D, Fisher DS. Fine-scale diversity and extensive recombination in a quasisexual bacterial population occupying a broad niche. Science. 2015;348: 1019–1023.

64. Lewis HC, Mølbak K, Reese C, Aarestrup FM, Selchau M, Sørum M, et al. Pigs as source of methicillin-resistant Staphylococcus aureus CC398 infections in humans, Denmark. Emerg Infect Dis. 2008;14: 1383–1389.

65. Ding T, Schloss PD. Dynamics and associations of microbial community types across the human body. Nature. 2014;509: 357–360.

66. Albrecht VS, Limbago BM, Moran GJ, Krishnadasan A, Gorwitz RJ, McDougal LK, et al. Staphylococcus aureus Colonization and Strain Type at Various Body Sites among Patients with a Closed Abscess and Uninfected Controls at U.S. Emergency Departments. J Clin Microbiol. 2015; doi:10.1128/JCM.01371-15

67. R Core Team. The R project for statistical computing. R Foundation for Statistical Computing web-site www.R-project.org Accessed June. 2014;9.

68. Lawson DJ, Hellenthal G, Myers S, Falush D. Inference of population structure using dense haplotype data. PLoS Genet. 2012;8: e1002453.

69. Darling AE, Mau B, Perna NT. progressiveMauve: multiple genome alignment with gene gain, loss and rearrangement. PLoS One. 2010;5: e11147.

70. Stamatakis A, Aberer AJ, Goll C, Smith SA, Berger SA, Izquierdo-Carrasco F. RAxML-Light: a tool for computing terabyte phylogenies. Bioinformatics. 2012;28: 2064–2066.

71. Yang Z. PAML 4: phylogenetic analysis by maximum likelihood. Mol Biol Evol. 2007;24: 1586–1591.

72. Li H, Durbin R. Fast and accurate short read alignment with Burrows-Wheeler transform. Bioinformatics. 2009;25: 1754–1760.

73. Méric G, Miragaia M, de Been M, Yahara K, Pascoe B, Mageiros L, et al. Ecological overlap and horizontal gene transfer in Staphylococcus aureus and Staphylococcus epidermidis. Genome Biol Evol. 2015; doi:10.1093/gbe/evv066

74. Human Microbiome Project Consortium. A framework for human microbiome research. Nature. 2012;486: 215–221.

